# Inference of population genetic structure from temporal samples of DNA

**DOI:** 10.1101/801324

**Authors:** Olivier François, Séverine Liégeois, Benjamin Demaille, Flora Jay

## Abstract

The recent years have seen a growing number of studies investigating evolutionary questions using ancient DNA techniques and temporal samples of DNA. To address these questions, one of the most frequently-used algorithm is based on principal component analysis (PCA). When PCA is applied to temporal samples, the sample dates are, however, ignored during analysis, which could lead to some misinterpretations of the results. Here we introduce a new factor analysis (FA) method for which individual scores are corrected for the effect of allele frequency drift through time. Based on a diffusion approximation, our approach approximates allele frequency drift in a random mating population by a Brownian process. Exact solutions for estimates of corrected factors are obtained, and a fast estimation algorithm is presented. We compared data representations obtained from the FA method with PCA and with PC projections in simulations of divergence and admixture scenarios. Then we applied FA with correction for temporal drift to study the evolution of hepatitis C virus in a patient infected by multiple strains, and to describe the population structure of ancient European samples.

## 1 Introduction

In recent years, the number of studies analyzing temporal samples of DNA or ancient DNA has increased dramatically, both for humans and for other organisms (Lazaridis *et al.*, 2014; Haak *et al.*, 2015; Mathieson *et al.*, 2015; Carroll *et al.*, 2015; Skoglund and Mathieson, 2018). In such studies, a central question concerns the inference of ancestral relationships between sampled populations (Slatkin, 2016). Evolutionary biologists and population geneticists have devised many methods for addressing this question. One of the most frequently-used method is based on principal component analysis (PCA) and projections of ancient samples on axes built from present-day samples (Patterson *et al.*, 2006, 2012). In population genetics, PCA is performed by finding the eigenvalues and eigenvectors, or axes, of the covariance matrix of allele frequencies. The highest order eigenvectors indicate the directions in the high dimensional allele-frequency space which account for most of the covariance. Individual samples are then plotted on the plane spanned by the first axes, offering a visual representation of the structure hidden in the data obtained with short computing time. Relative distances in the reduced space indicate their similarity and their ancestral relationships (McVean, 2009). When PCA or PC projections are applied to analyze temporal samples, information on sample dates is, however, usually omitted in the computation of eigenvalues and eigenvectors (Slatkin, 2016; Slatkin and Racimo, 2016; Harris and DeGiorgio, 2017).

Previous studies have reported that time differences in samples are reflected in the principal axes of a PCA (Skoglund *et al.*, 2014), creating sinusoidal shapes similar to those observed with geographic samples (Novembre and Stephens, 2008). The combination of both time and spatial heterogeneity in sampling further modify the patterns observed in PCA. Local dispersal through time causes ancient samples to be shrunk toward the center of the PC plot and not to cluster with their present-day counterpart despite no major discontinuity in the demographic process (Duforet-Frebourg and Slatkin, 2016). Sinusoidal distortions linked to gradients and longitudinal data also occur in various fields, and are called *horseshoes* or *arches*. Those distortions complicate the interpretation of multidimensional scaling, local kernel methods and ordination analysis (Hill and Gauch, 1980; Diaconis *et al.*, 2008). Supervised methods that combine ancient and modern samples by using PC projections on present-day samples also suffer from some statistical issues. PC projections exhibit a shrinkage bias toward the center of the principal axes, and this bias could increase in analyses of temporal samples (Lee *et al.*, 2010). Since those biases could lead to misinterpretations or to incorrect estimates of individual ancestry, it is important to propose methods that correct principal components when temporal samples are analyzed for descriptive purposes.

Corrections of sinusoidal patterns arising in principal components have been proposed when distortions are caused by spatial auto-correlation in geographic samples (Frichot *et al.*, 2012). Similarly, Kalaitzis and Lawrence (2012) have proposed to remove temporal correlations leaving residual variance with residual component analysis. Modified versions of the STRUCTURE algorithm – which is closely related to PCA – were also developed to integrate corrections based on spatial or temporal diffusion models (Pritchard *et al.*, 2000; Caye *et al.*, 2018; Joseph and Pe’er, 2018). In this study, we introduce a new factor analysis (FA) method for visualizing hidden structure and for describing ancestral relationships among samples collected at distinct time points in the past. Based on a diffusion approximation, our approach approximates allele frequency drift in a random mating population by a Brownian process. Using the Karhunen-Loève theorem, we propose a representation of the factor model in which additional covariates, representing temporal eigenvectors, are introduced in the model. Our model assumes informative Gaussian prior distributions for the effect sizes of the temporal covariates. Exact solutions for time-corrected factors are obtained, and a fast algorithm based on singular value decomposition (SVD) is proposed. We compare corrections for temporal drift in FA with PCA in coalescent and generative simulations of divergence and admixture scenarios. We eventually apply corrections for temporal drift to study the evolution of hepatitis C virus in a patient infected by multiple viral strains, and to describe population structure for DNA samples from ancient Europeans and Eurasians.

## 2 New Method

This section introduces a new factor analysis method for describing ancestry among samples taken at distinct time points in the past. The objective is to propose a factorial decomposition of the data matrix similar to a PCA, in which the individual scores are corrected for the effect of allele frequency drift through time. The scores, called factors, will be obtained as maximum-a-posteriori estimates in a Bayesian model.

### Model

Let **Y** be an *n × p* matrix of genotypic data, where *n* is the number of individual samples and *p* is the number of markers, typically represented as single nucleotide polymorphisms (SNPs). We suppose that the data are centered, so that the mean value for each column (or marker) is null. We also suppose that each sample, *i*, is associated with a sampling date, *t*_*i*_, corresponding to the age of the sample. The dates are normalized to span the unit interval 0 < *t*_1_ ≤ ⋯ ≤ *t*_*n*_ ≤ 1. Here, time has a forward representation. The date *t*_1_ corresponds to the most ancient sample, and *t* = 1 represents samples at present time. Our FA model takes the following form

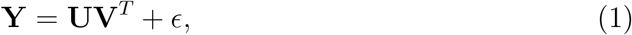

where **U** is an *n × K* matrix of scores, **V**^*T*^ is a *K × p* matrix of loadings. The number of factors, *K*, can be set to any number smaller that *n* and *p* depending on how drastically one wants to reduce the dimension of the data (and approximate the data matrix). It can be set to the number of ancestral groups minus one when this information is known. The individual scores contained in the *K* column vectors, **u**_1_, …, **u**_*K*_, of the matrix **U** reflect the ancestral relationships among samples (Patterson *et al.*, 2006). To incorporate corrections for temporal drift, we model the error term, *ϵ*, as follows

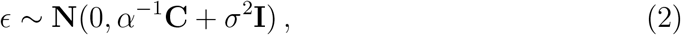

where **N**(0, *α*^−1^**C** + *σ*^2^**I**) is the multidimensional Gaussian distribution with mean 0 and covariance matrix *α*^−1^**C** + *σ*^2^**I**, *α* is a precision (scale) parameter for temporal drift, *σ*^2^ is the variance of the residual error, and **I** is the *n × n* identity matrix. We suppose that **C** is an *n × n* covariance matrix given by

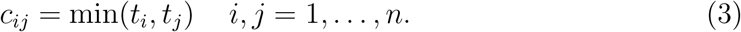

The definition of the covariance matrix, **C**, is related to the covariance function of the Brownian process. This model assumption corresponds to the diffusion approximation of allele frequency drift in a random mating population conditional on non-fixation of alleles in the population (Kimura, 1964, 1983). The diffusion approximation underlies the development of several recent methods of ancestry estimation similar to our model (Patterson *et al.*, 2012; Pickrell and Pritchard, 2012; Peter, 2016; Joseph and Pe’er, 2018). As a consequence of the definition, the variance of allele frequencies is proportional to time. In applications, we normalized the sample dates so that *t*_1_ corresponds to the variance of allele frequencies in the oldest sample.

### Factor estimates

To compute the factor matrix, **U**, in the model equation (1), we turned to an equivalent formulation of this equation

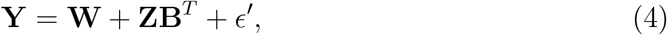

where the residual noise is described by

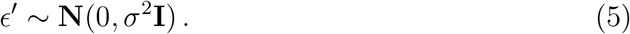

In this formula, effect sizes, *B*_*j*_, *j* = 1, ⋯, *p*, are introduced, and considered as i.i.d. random variables with univariate Gaussian prior distribution *N* (0, *α*^−1^). A latent matrix, **W** = **UV**^*T*^, has a non-informative prior distribution. After a spectral decomposition of the covariance matrix **C**, we define

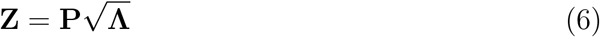

where **P** is a unitary matrix of eigenvectors, and **Λ** is the diagonal matrix containing the eigenvalues of **C**

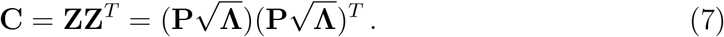

Based on the Karhunen-Loève theorem (Loève, 1948), the diagonal terms of **Λ** can be approximated as

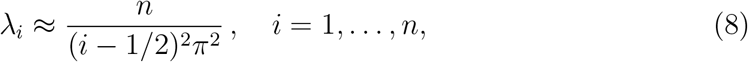

and we have

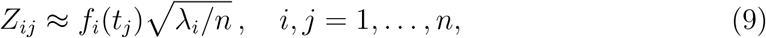

where *f*_*i*_(*t*) is defined as 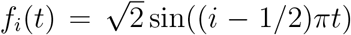 for all *t* in the interval [0, 1]. According to these results, the eigenvectors of the covariance matrix have sinusoidal shapes, and a diffusion model is consistent with the arch effect observed in principal components of genetic variation (Skoglund *et al.*, 2014).

Statistical estimates of the matrices **U**, **V** and **B** can be obtained by maximizing a posterior distribution in a Bayesian framework. This approach amounts to finding the minimum of the following loss function

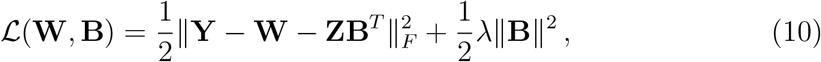

where we have set *λ* equal to the inverse of the temporal signal-to-noise ratio, *λ* = *ασ*^2^. Finding the matrices **W** and **B** that minimize the loss function 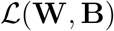 is equivalent to computing their estimates in a latent factor regression model with ridge penalty (Frichot *et al.*, 2013). According to Caye *et al.* (2019), the latent matrix, **W**, minimizes the following loss function

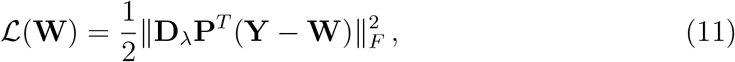

where **D**_*λ*_ is a diagonal matrix with coefficients equal to

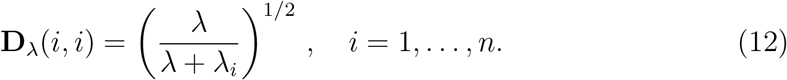

The estimate of **W** is provided by the best approximation of rank *K* of the matrix **Y**, where “best approximation” is related to the following matrix norm

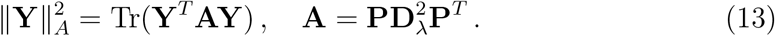

In closed form, the optimal solution is equal to

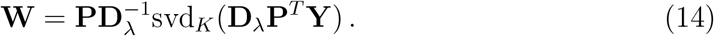

The *K* corrected factors forming **U** and their associated loadings, **V**, can be obtained from the SVD of the matrix **W** (see Table S1). For very large data sets, a modification of the SVD based on random projections could provide an accelerated version of the algorithm (Halko *et al.*, 2011).

### Software availability

A short working `R` code presenting the algorithm in a selfcontained way is provided in Table S1. The method described in this section is currently implemented as an `R` package `temporalFA`.

## 3 Results

### Horseshoe effect

To provide an example of distortion arising in PCA due to uncorrected temporal drift, we performed a simulation of a coalescent model for forty-one samples with ages ranging from 0 to 4,000 generations in a population with effective size *N*_e_ = 10, 000. The sample dates in the simulation corresponded to an interval of 100 generations. Covariance among samples was smaller for the most ancient samples than for the most recent samples, and it increased linearly with time (Figure 1A). The patterns observed in the sample covariance matrix were highly similar to those obtained in a theoretical covariance function corresponding to a Brownian process (Figure 1C). The PC plots of individual samples exhibited sinusoidal patterns, in which the most ancient and recent samples were placed at both extremes of a horseshoe (Figure 1B). Correcting for temporal drift, the factor analysis plot displayed a single cluster grouping all samples without any apparent structure among samples (Figure 1D). This last result showed that distortion due to temporal drift was correctly removed in a factor analysis using a Brownian model of genetic drift.

**Figure 1.**
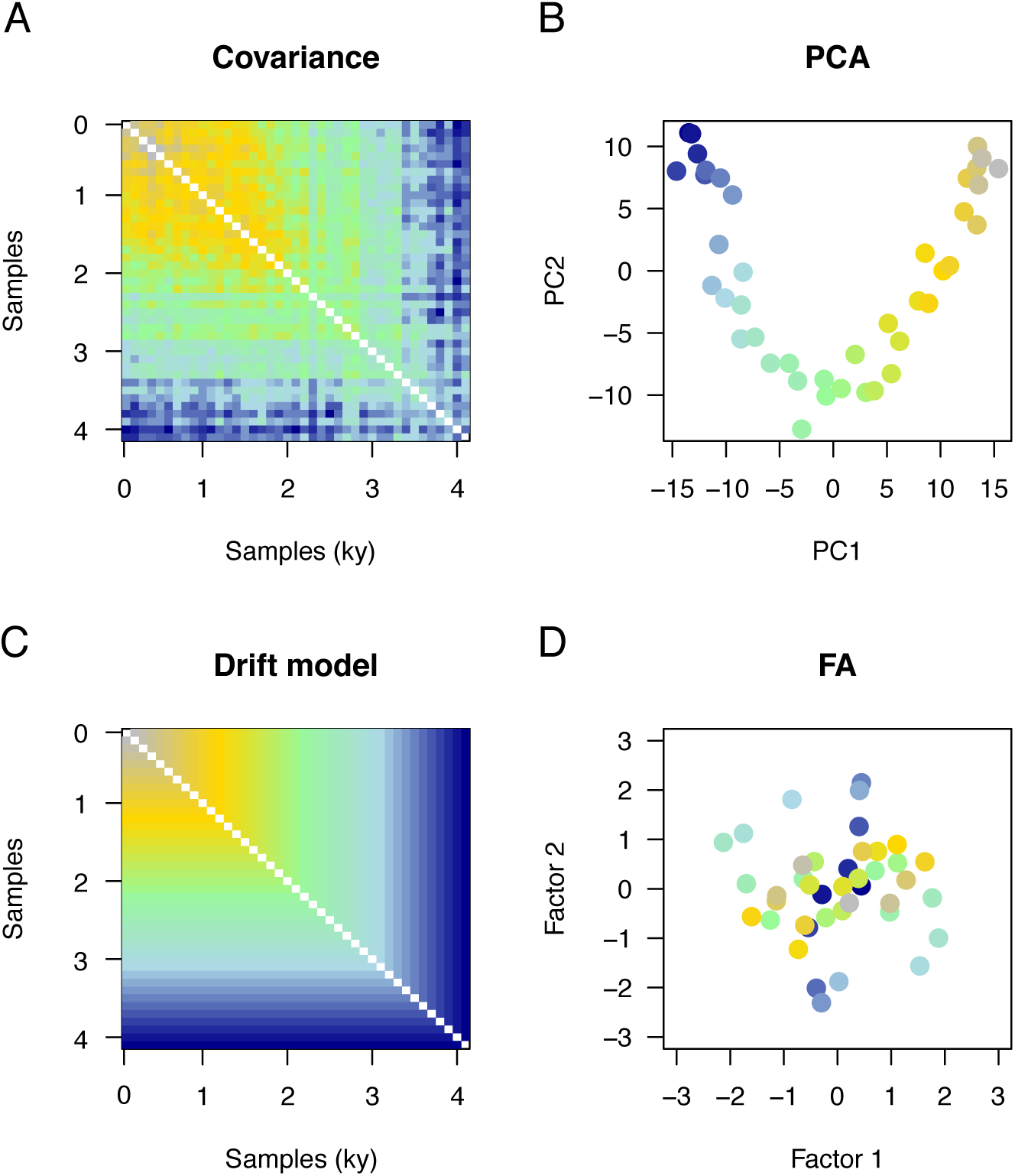
Horseshoe effect. Coalescent simulation of allele frequencies drifting through time in a single population (*N*_e_ = 10, 000). Forty-one samples with ages ranging from 0 (present, grey color) to 4,000 generations (past, dark blue color) were simulated. A) Covariance matrix for observed samples B) PC plot for individual samples, C) Brownian covariance matrix used as a correction model, D) Factor analysis plot showing correction for temporal drift. In covariance matrices, the blue color indicates lower values whereas the yellow and grey colors indicate higher values.

### Divergence model

In a second series of experiments, we simulated models of divergence of two populations. In coalescent simulations, twenty-four samples with ages ranging from 0 to 1,000 generations were simulated, corresponding to a sampling interval of 100 generations and four present-day individuals. In a PCA of simulated samples, PC1 reflected the level of divergence between populations while PC2 represented temporal drift (Figure 2A). Correcting for temporal drift, the factor analysis plot exhibited two clusters without any apparent structure within each group (Figure 2B). The Davies-Bouldin clustering index reached higher values in the FA plots than in the PC plots, meaning that the clusters were better characterized and better represented populations of origin in FA than in PCA (Figure 3C). In generative model simulations, factor 1 in FA better explained the hidden factor than did the first PC in PCA (Figure 3D). The results provided evidence that correcting for temporal drift in FA revealed population structure hidden in the noisy data.

**Figure 2.**
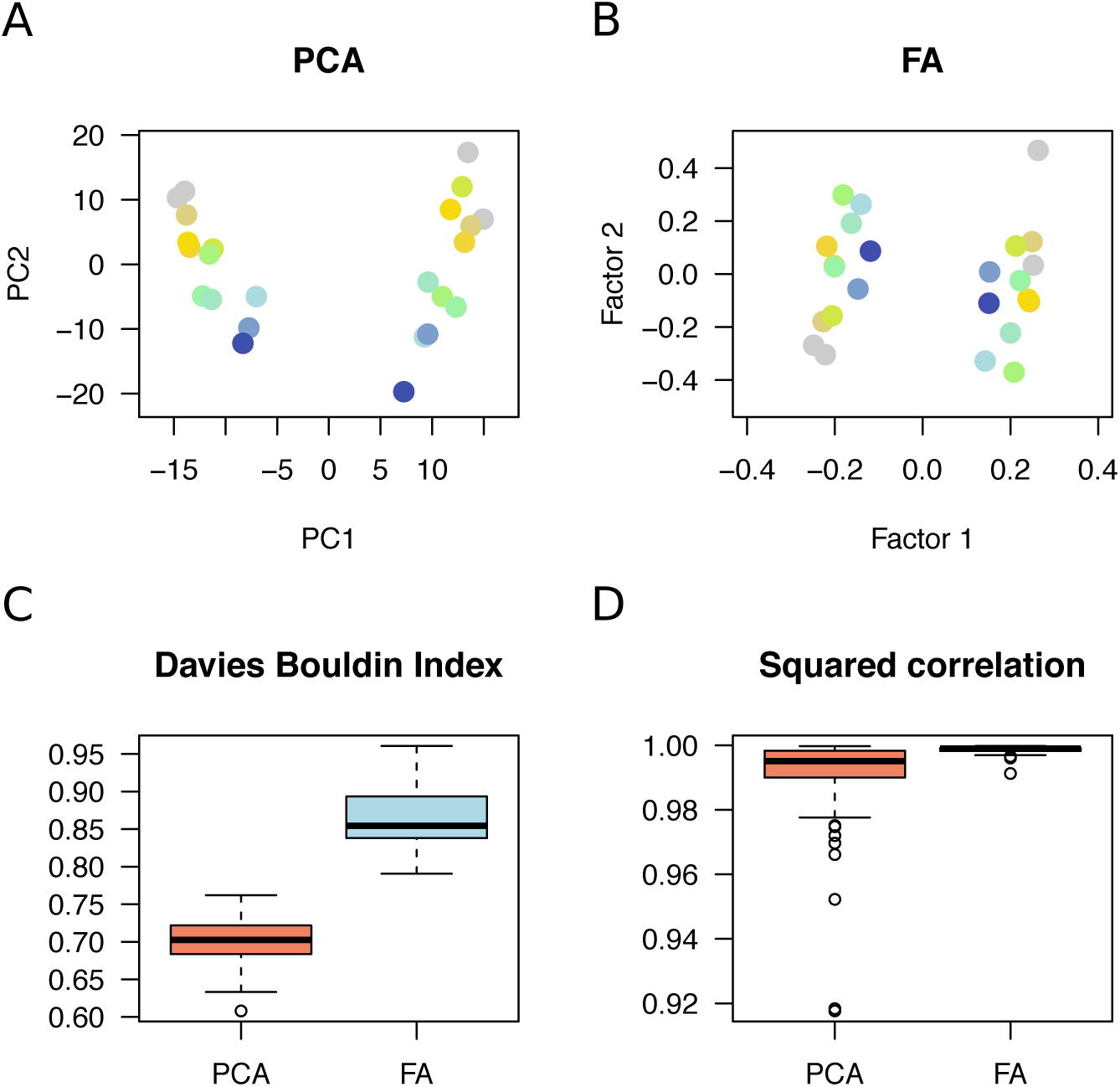
Simulation of two-population models. Twenty-four samples with ages ranging from 0 (present, grey color) to 1,000 generations (past, dark blue color) were simulated. A) Typical PC plot for observed samples, B) Factor analysis plot showing correction for temporal drift, C) Clustering index for PCA and FA results (100 coalescent simulations), D) Squared correlation between PC1 - Factor 1 and a true factor having two modes (100 generative model simulations).

**Figure 3.**
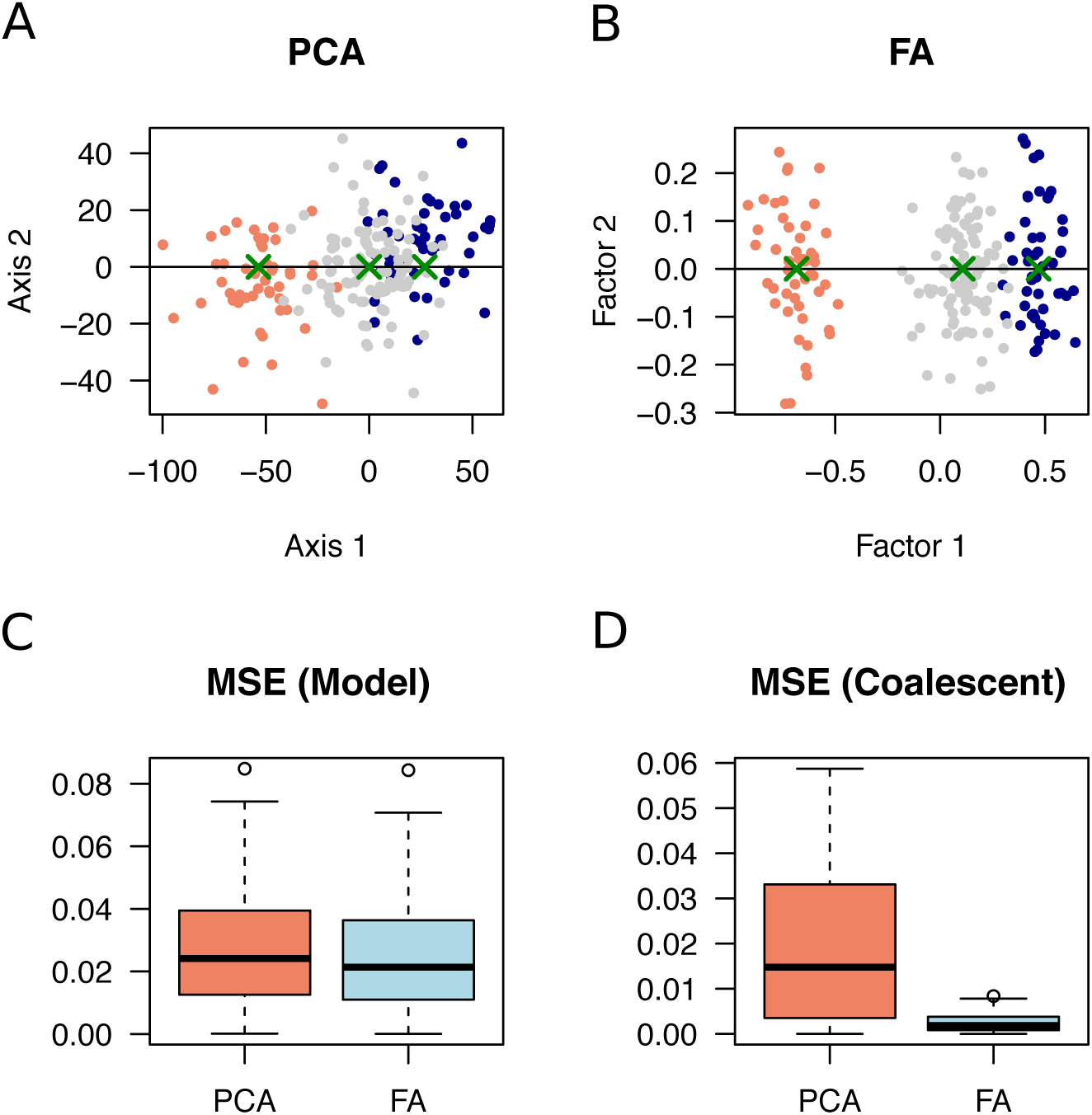
Simulation of admixture models. One hundred samples were simulated for present-day admixed individuals (admixture rate 25 and 75%, grey color) and samples from two ancestral populations (age 1,000 generations, orange and blue colors). A) Typical plot for PC projection of ancient samples onto the admixed population showing a shrinkage effect, B) Factor analysis plot showing correction for shrinkage, C) Mean square error for estimates of admixture proportions from PC projections and FA plots (100 generative model simulations), D) Mean squared error for estimates of admixture proportions (100 coalescent simulations). Green crosses represent population centers, from which admixture estimates were computed.

### Admixture models

In another series of experiments, we considered admixture models in which an ancestral population splits into two sister populations 1,300 generations ago. The two divergent populations came into contact 500 generations ago, giving rise to descendants having 75% ancestry in the first ancestral population and 25% ancestry in the second ancestral population. One hundred present-day individuals were sampled from the admixed population, and fifty individuals were sampled from each ancestral population before the admixture event (1,000 generations ago). Artificial genotypes generated according to a Brownian model were also used to simulate levels of admixture similar to those observed in coalescent models (see Methods). The objective of the experiments was to compare the results of PC projections of ancient samples onto the present-day population with those obtained in factor analysis with correction for temporal drift. Typical plots for PC projections exhibited a shrinkage effect in which the projected samples were shifted toward zero, and closer to the admixed population than expected (Figure 3A). The shrinkage effect was even more pronounced in coalescent simulations than in generative model simulations (Figure S1 and Figure 3C-D). Correction for temporal drift in factor analysis removed the shrinkage effect, and, in the FA plot, the locations of centers of ancestral clusters reflected admixture levels more precisely than in PC plots (Figure 3B). The mean squared errors for estimates of admixture proportions were higher in PC projections than in FA plots both in generative and in coalescent simulations (Figure 3C-D). The results showed that correcting for temporal drift in FA improved the representation of admixed individuals and their source populations compared with projections on present-day individual PCs.

### Hepatitis C virus infection

To follow chronic infection in a non-responder hepatitis C patient treated in the 2000’s, we studied *n* = 1, 934 samples of viral RNA sequences over a period of thirteen years (Caporossi *et al.*, 2019). The patient was coinfected by viral strains from two HCV genotypes, 4k and 1b. Height serum samples were available from years 2002 to 2014. Treatment with dual therapy had been administered for six months after the beginning of the follow-up period. A PC plot of viral samples displayed a pattern similar to those observed in simulations of divergence models (Figure 4A and Figure 2A). PC1 reflected divergence among the samples classified in distinct viral types, and PC2 was influenced by the ages of the samples. After correction for temporal drift in a FA plot, viral particles were grouped according to their phylogenetic classification (Figure 4B). In the FA plot, a first cluster consisted of 1b strains from year 2003 to 2014. A second cluster consisting of 4k strains exhibited some degree of substructure, separating samples taken during treatment (year 2003) to the other samples. An interpretation of this result was that 1b strains had mainly evolved through drift after treatment, whereas 4k strains might had experienced other evolutionary changes, suggesting selection on this genotype during the evolution of the disease (Caporossi *et al.*, 2019).

**Figure 4.**
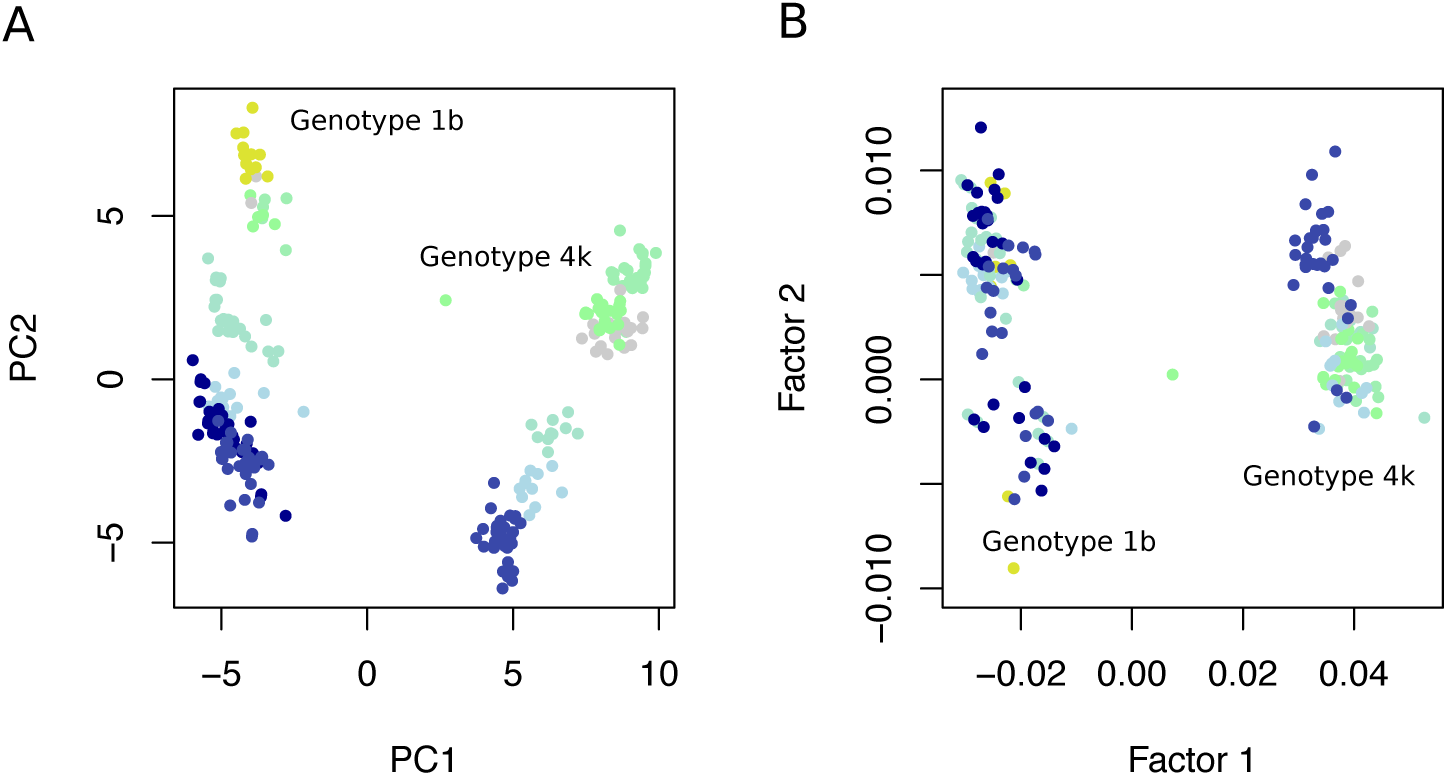
Hepatitis C virus infection. Longitudinal study of a single non-responder patient infected by two viral strains (HCV genotypes 1b and 4k). A total of *n* = 1, 934 viral samples were collected from years 2002 to 2014. Dark blue color corresponds to the oldest samples, while yellow and grey colors correspond to the most recent samples. A) PC plot of viral samples, B) factor analysis with correction for temporal drift. A few outlier individuals detected in the FA plot are not shown in the plot.

### Ancient European genomes

We used PC projections and a Brownian model of factor analysis to study a merged data set consisting of 155k SNP genotypes for 249 present-day European individuals and 386 ancient samples from Eurasia. The ages of ancient individuals were less than 12,080 years cal BP, and individuals were selected to be close to present-day Europeans in a preliminary FA analysis to leverage the effect of low genomic coverage on factor one. The data set contained ancient samples mainly from (Olalde *et al.*, 2018; Mathieson *et al.*, 2015; Haak *et al.*, 2015; Mathieson *et al.*, 2018). First, we computed principal components on present-day samples, and projected the ancient samples on the first two PCs (Figure S2). We also computed factors with temporal correction for present-day and ancient samples, choosing the hyper-parameter so that the factors correlate with principal components on present-day individuals (*λ* = 2 × 10^−6^, Multiple *R*^2^ = 0.97 for factor 1, *R*^2^ = 0.75 for factor 2, *P* < 10^−10^, Figure S3). Both analyses revealed a similar pattern, in which most ancient samples from Ukraine and all samples from Scandinavia, including hunter-gatherers from Latvia, were close to present-day Finnish samples, ancient samples from Great Britain were close to present-day British samples, and ancient samples from Anatolia and Israel were close to present-day southern Europeans. Ancient samples from Iran, Armenia and Iraq formed a distinct group.

Next, we performed an unsupervised time-corrected factor analysis considering ancient samples only. In this analysis, sample ages explained 0.2% of the variance in factor 1 and 5.4% of the variance in factor 2, showing that temporal bias was correctly removed from the first two factors (*λ* = 10^−3^). The FA plot exhibited four main clusters and a pattern of variation strongly consistent with the geographic origin of samples (Figure 5, see Figure S4 for a definition of clusters). A first cluster grouped ancient samples from Ukraine, Latvia and Sweden (Figure 5, green color). Ages in the Scandinavian cluster 1 were around 7,671 years BP (mean value, SD = 1,710 years). A second cluster grouped ancient samples from Russia, including samples from Samara of the Yamnaya culture, Central Europe and Great Britain (Figure 5, dark blue stars to golden points). Ages in cluster 2 were around 3,832 years cal BP (SD = 1,570 years). A third cluster grouped individuals from Anatolia and Israel, Southern Europe and Great Britain (Figure 5, brown stars to golden points). Ages in the southeastern cluster 3 were around 6,079 years cal BP (SD = 1,311 years). A fourth cluster grouped samples from Central Asia (salmon triangles). Samples from Bronze age Great Britain (4,300 years BP) were grouped in cluster 2, whereas samples from the neolithic period and from the same region were found in cluster 3 representing Southeastern Europe. More generally, samples with ages older than around 4,500 years BP were grouped in the southeastern cluster, while more recent samples from the bronze age clustered with ancient North Eurasian, Russian and Steppe samples in the central cluster 2. Discontinuities in ancestry reflected in factor 1 were observed for samples from Great Britain, Germany and Hungary (Figure 6). In Hungarian samples, a linear trend was observed for the period 4,500 - 8,000 years BP, consistent with levels of hunter-gatherer ancestry detected in (Lipson *et al.*, 2017).

**Figure 5.**
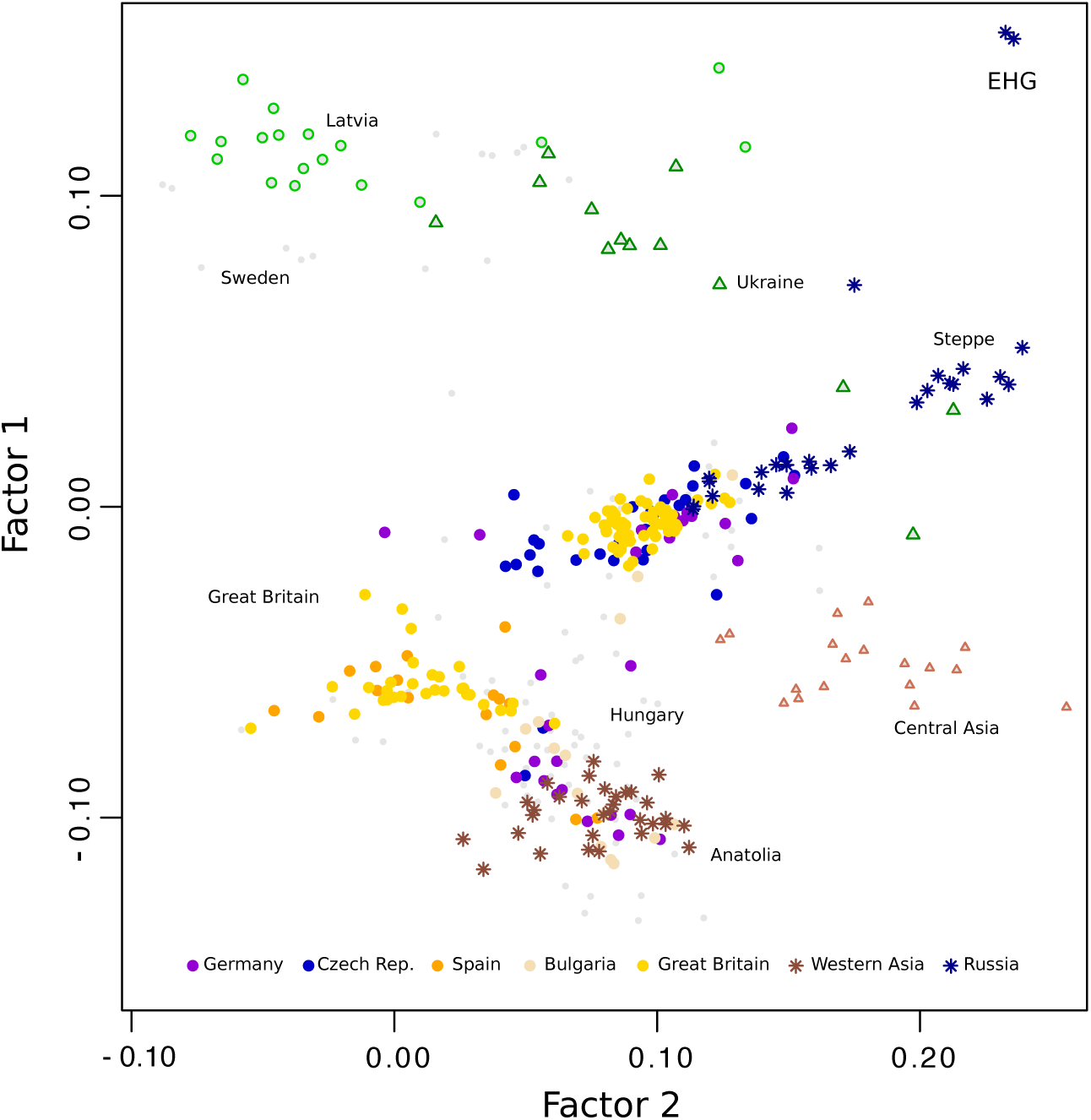
Ancient European genomes. Factor analysis of 386 ancient Eurasian individuals with ages ranging between 400 and 12,000 years BP. Four main groups represent individuals from 1) Northern Europe and Ukraine (green color), 2) Russia, Steppe, Central Europe and the British Isles (average dates around 4k years BP, blue color), 2) Near East, Southern Europe and the British Isles (average dates around 6k years BP, brown color), and 4) Central Asia (Salmon color).

**Figure 6.**
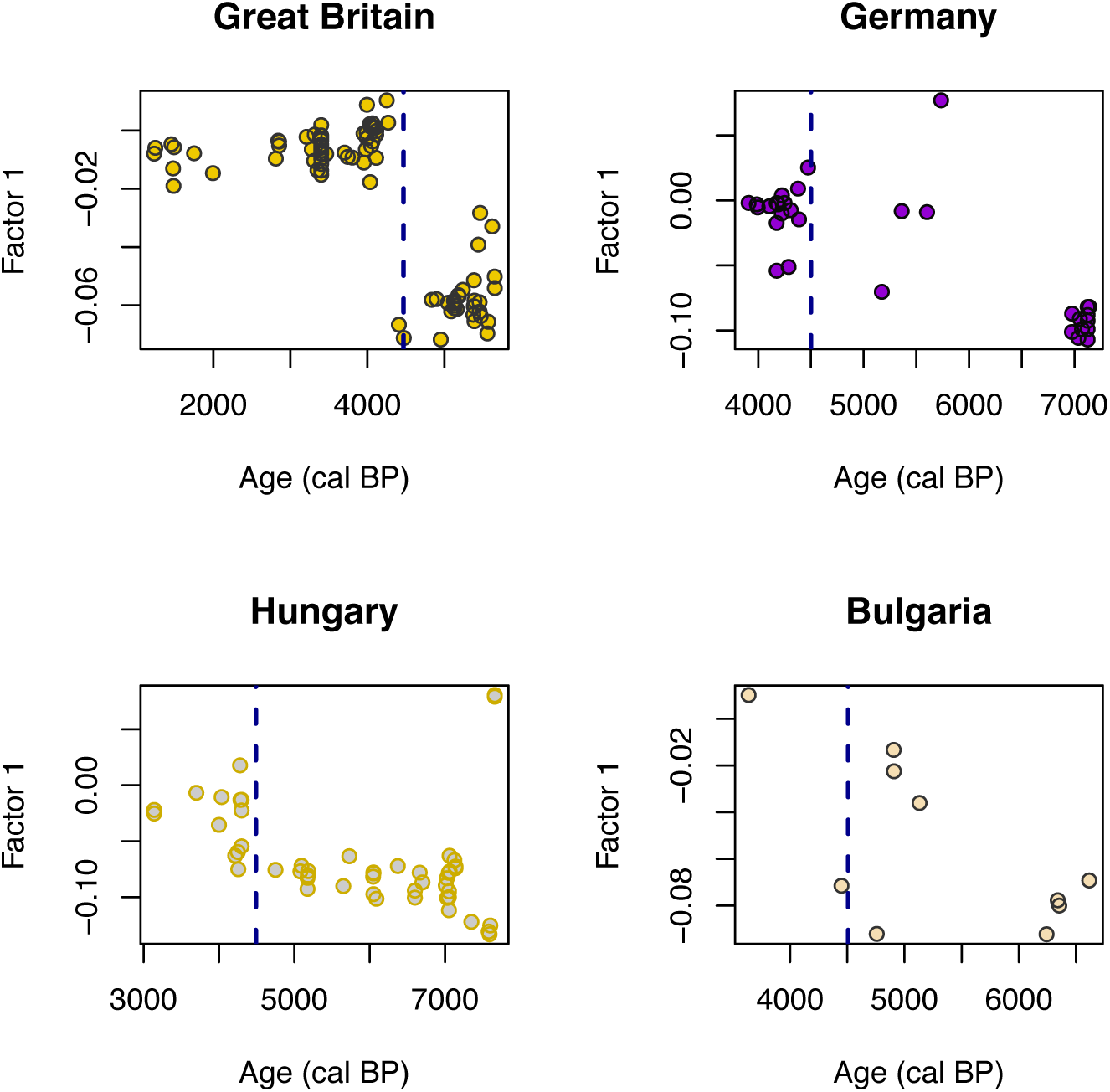
Factor 1 as a function of age (years cal BP). Factor 1 of a temporal FA displayed as a function of age for samples from Great Britain, Germany, Hungary and Bulgaria. The data support a major change in genetic mixture of individuals from Great Britain, Germany, Hungary around 4,500 years BP (dashed line).

To assess the genetic ancestry of samples from Great Britain, a second FA was performed. This analysis isolated British samples from ancient North Eurasians and ancient Near Easterners, considered as putative source populations (Figure 7). British samples with dates earlier than 4,300 years cal BP clustered with samples from the Near East. Samples with dates around 4,300 years cal BP (early bronze age) were close to samples from Russia, and a genetic discontinuity was observed with more ancient samples from Anatolia. Estimating admixture coefficients from factor 1, the early bronze age samples shared around 64% of their ancestry with the North Eurasian samples and 36% with the Neolithic Easterners. Samples from the middle bronze age (around 3,300 BP) formed a distinct group, suggesting a more complex history than two waves of invasions in the British Isles.

**Figure 7.**
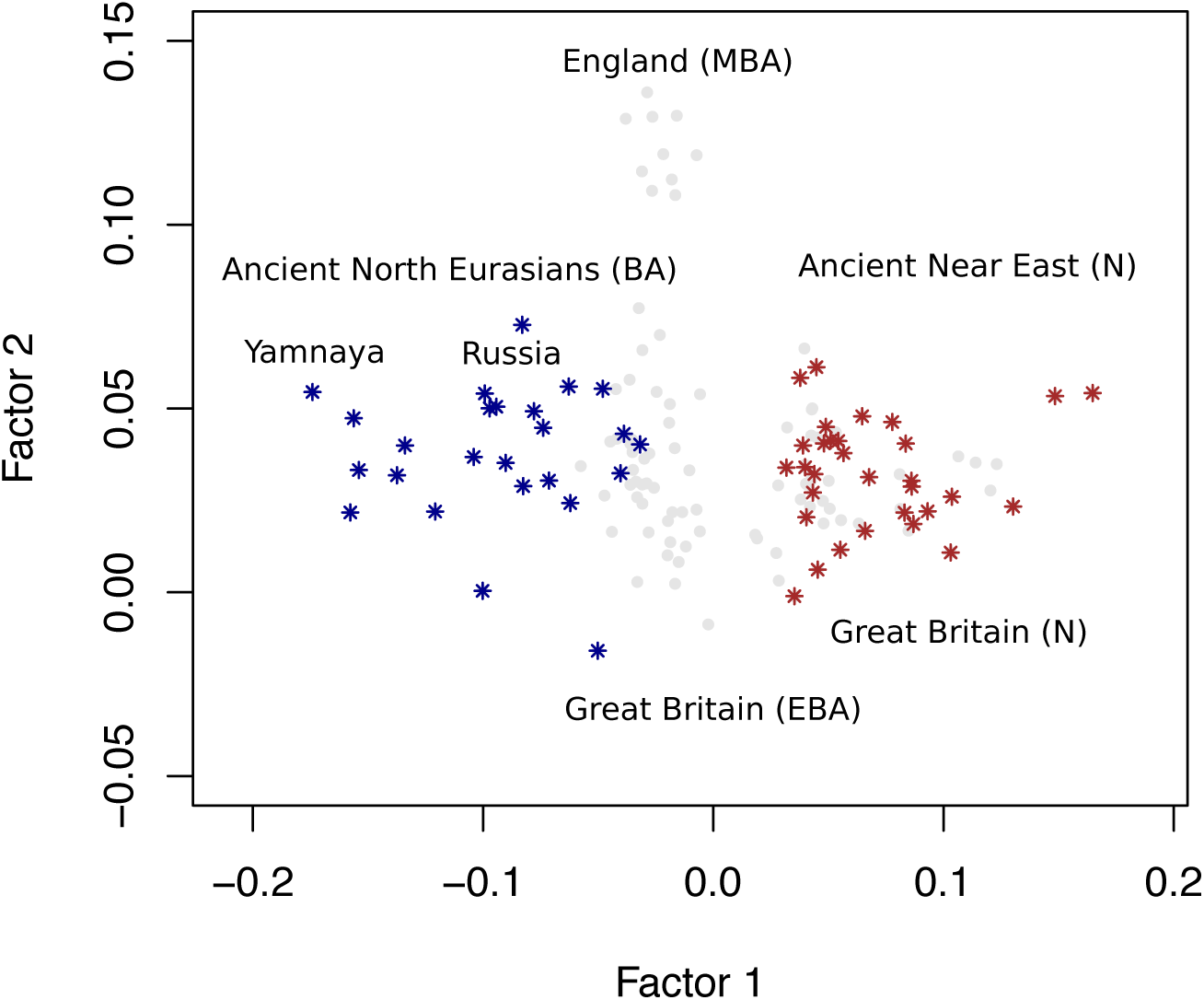
Ancestry of ancient samples from Great Britain. FA plot for samples from ancient North Eurasians (Russia and Samara with Yamnaya Culture, dark blue color), from ancient Near East (Neolithic Anatolia and Israel, brown color), and from Great Britain (grey color). The samples from Great Britain cluster in three groups: Neolithic (N), Early Bronze Age (EBA), Middle Bronze Age (MBA).

Finally, a larger set of 697 ancient samples was considered for replication of PC projection and unsupervised FA results. PCA and FA plots yielded similar descriptions of the data when a larger set of ancient samples was considered (Figure S5–S6).

## 4 Discussion

We introduced a new factor analysis method for describing ancestral relationships among DNA samples collected at distinct time points in the past. Like in PCA, the method is based on a factorial decomposition of the data matrix into a product of score and loading matrices. The most important difference with the PCA approach is that individual scores in FA were corrected for the effect of temporal drift in allele frequency. Based on a diffusion approximation, we approximated allele frequency drift by a Brownian process, and an efficient algorithm based on the singular value decomposition computed the factor estimates.

Using a Brownian model of genetic drift, we compared the results of FA with those of PCA and PC projections in simulations of divergence and admixture. In divergence scenarios, distortions due to temporal drift were removed in FA. Correcting for temporal drift revealed hidden population structure better than did a PCA. In admixture scenarios, estimates of ancestry coefficients were more accurate in FA than those inferred from principal components. In those simulations, correcting for temporal drift allowed a better representation of admixed individuals than PC projections.

Next, we applied temporal corrections to study the evolution of hepatitis C virus in a patient infected by multiple strains. After correction for temporal drift, viral strains clustered according to their phylogenetic classification. In agreement with the fact that the patient did not respond to treatment, FA suggested that 1b strains had mainly evolved through drift after treatment. Evidence for substructure within 4k samples suggested an action for other evolutionary processes among those strains. Caporossi *et al.* (2019) reported that nucleotide diversity was higher in 1b time samples than in 4k time samples, which might indicate that drift was more important in the 4k population. With the FA result, this suggests that distinct corrections should be applied to 4k and 1b samples. We performed a separate FA with 4k samples only (not shown), and the observed substructure persisted. Overall, the FA plot supported the hypothesis that drift was not the only process acting on the genetic diversity of 4k genotypes, and that those strains might have experienced some form of selection during the course of disease evolution Caporossi *et al.* (2019).

In a re-analysis of a merged data set of ancient DNA filtering out SNPs with high levels of missing data and genomes of low coverage, we implemented correction for temporal drift to describe ancestry in samples from ancient Europeans and Eurasians. After correction, the patterns observed in FA plots were consistent with those observed in projections of ancient samples on axes built on the 1,000 Genomes data. The factor analysis supported the hypothesis that a major change in genetic mixture of individuals occurred in Great Britain and in continental populations around 4,300 years BP (Olalde *et al.*, 2018). Observed FA patterns were more consistent with geography in than those in PC projections, suggesting a role of localized gene flow unseen in previous analyses at the continental scale. Our analysis provided a visual representation of Bronze age British samples consistent with the proportion of North Eurasian and steppe ancestry of the original (Olalde *et al.*, 2018).

In conclusion, including corrections for temporal drift resulted in an algorithm with a computational cost similar to a PCA. Determining the model hyper-parameter was based on simple approaches, computing a correlation between sample dates and first FA scores. Our study showed that the FA method corrected biases observed in PC plots successfully. A useful and important feature of the new approach was to avoid supervised analyses in which unbalanced samples over-representing present-day individuals are utilized. The unsupervised approach based on FA revealed details of population structure masked in PC projections, and was generally more accurate than principal component analysis of population structure for ancient samples.

## 5 Materials and Methods

### Coalescent simulations

We used the computer program *msprime* to simulate temporal samples for individuals at distinct time points in the past (Kelleher *et al.*, 2016). Firstly, a single population of *N_e_* = 10, 000 individuals was simulated during 4,000 generations. An individual was sampled every 100 generations, resulting in 41 samples with ages ranging between 0 (present-day) and 4,000 generations. A total of around 9,000 SNPs were simulated for each individual. Secondly, a divergence model was considered in which an ancestral population of effective size *N_e_* = 10, 000 split into two sister populations of equal sizes 1,500 generations ago. Twenty-four individuals with ages ranging from 0 to 1000 generations were sampled every 100 generations (four present-day individuals were simulated), and around 8,800 SNPs were simulated for each individual. One hundred replicate data sets were created with the same demographic parameters. For each simulation, the Davies-Bouldin index was computed (Davies and Bouldin, 1979). The Davies-Bouldin index is a metric for evaluating the degree of clustering in multidimensional data, and ranges between zero and one. Corrections for temporal drift in allele frequency are expected to provide index values closer to one than those for principal components. Thirdly, an admixture model was considered in which an ancestral population of effective size *N_e_* = 10, 000 split into two sister populations of equal sizes 1,300 generations ago. The two divergent populations came into contact 500 generations ago, and this event gave rise to a third population. Individuals in the admixed population shared 75% ancestry with the first ancestral population, and 25% ancestry with the second ancestral population. One hundred individuals were sampled from the admixed present-day population, and fifty individuals were sampled from each ancestral population, 1,000 generations ago. A total of around 9,600 SNPs were simulated for each individual. One hundred replicate data sets were created with the same demographic parameters. For each simulation, we computed the centers of the ancestral and admixed population on the first axis, and we estimated admixture proportion based on the ratio of distances between population centers. We also did this for the first factor with correction for temporal drift. We eventually computed mean squared estimation errors both for PCA and for FA estimates.

### Generative model simulations

Since the correction method is not restricted application to ancient DNA, we performed a series of experiments using the generative model defined in equation (1)

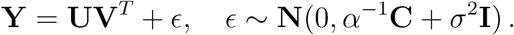

The objective was to evaluate statistical errors for latent factor estimates in a general context. The generative model simulations have the advantage of creating artificial data for which the ground truth is available. Based on a genealogical interpretation of principal components, we devised two series of simulations (McVean, 2009). The first scenario considered a divergence model in which two populations evolved without gene flow. In this case, populations were grouped separately along the first factor, and their divergence time was represented by the distance separating the group means.

The samples were taken at random times in the past and correlated noise was included in the data matrix. The second scenario considered an admixture model in which two populations diverged in the distant past and an admixture event occurred recently. Half of the samples were ancient, taken from the ancestral populations at random times in the past, and the other half of the samples were collected from the admixed population in present time.

For the divergence model, the factor matrix **U** contained *K* = 3 factors, simulated as Gaussian independent random variables. The standard deviation for first factor, *s*_1_, measured divergence between the two ancestral populations, and was varied in the range from to 2 to 10. Factors 2 and 3 had lower standard deviations, respectively equal to *s*_2_ = 1.5 and *s*_3_ = 0.5, so that **u**_1_ contained the largest genomic information. The λ parameter, representing an inverse temporal signal-to-noise ratio, was chosen in the range [10^−1^, 10^−6^]. The number of samples, *n*, was equal to 200, and the number of markers was kept to *p* = 1, 000. Loadings were simulated as independent standard Gaussian random variables, and the residual variance was set to *σ*^2^ = 1. For each simulation the squared correlation between the true **u**_1_ and estimated factor **û**_1_ was computed.

For the admixture model, the factor matrix **U** contained *K* = 3 factors. In the first factor, the two ancestral populations were positioned (with a standard deviation of 1) so that the distance separating their centers, *d*_1_, measuring divergence between them, was varied in the interval [10,12]. Factors 2 and 3 had standard deviations equal to *s*_2_ = 1.2 and *s*_3_ = 1. Admixed individuals were positioned so that center was at relative distance *a* from ancestral population 1, and 1−*a* from ancestral population 2, where *a* represents the ancestry contribution of population 1 to modern samples. The simulated ancestry coefficients ranged between *a* = 0.2 and *a* = 0.4. The *λ* parameter was set to *λ* = 5.10^−2^. The number of samples was set to *n* = 200, and the number of markers was kept to *p* = 5, 000. The loadings were simulated as independent Gaussian, N(0,0.2), random variables, and the residual error was set to *σ* = 0.1. We performed a total of 100 simulations. For each simulation, the squared correlation between the true **u**_1_ and estimated factor **û**_1_ was computed, and an estimate of the ancestry coefficient was provided, based on the relative positions of cluster means in **û**_1_.

### Hepatitis C virus data

To understand chronic infection in non-responder hepatitis C virus (HCV) patients treated with dual therapy in the 2000’s, Caporossi *et al.* (2019) performed deep sequencing on the NS5B (381 bp) region of the viral genome for a patient followed at Grenoble-Alpes University Hospital. The patient had a known date of infection because of an identified transmission event due to transfusion. The patient was treated with dual therapies based on pegylated interferon and ribavirin. The treatment had been administered for six months from January to June 2003, and a total of height serum samples were available for a follow-up period of 13 years. Co-infection by viral genotypes 4k and 1b was detected, and *n* = 1, 934 RNA samples from years 2002 to 2014 were studied.

### Ancient Human DNA samples

A merged data set consisting of genotypes for 1,820 ancient and present-day individuals compiled from published papers was downloaded from David Reich’s repository (https://reich.hms.harvard.edu/). The downloaded data matrix contained up to 1.23 million positions in the genome. Considering age defined as average of 95.4% date with range in cal BP computed as 1950 CE, Eurasian samples with age less than 12,080 years were retained. The data matrix was filtered out for samples falling far outside of the present-day Europeans in a preliminary FA analysis, leading to a median genomic coverage of 3.35x and a minimum coverage of 0.51x in the final data set. Only genomic positions with less than 25% of missing genotypes were analyzed. Missing genotypes were imputed by using a matrix completion algorithm based on sparse non-negative matrix factorization (Frichot and François, 2015; Frichot *et al.*, 2014).

The resulting data set contained 155,682 genotypes for 249 present-day European individuals from the 1,000 Genomes project (phase 3) and 386 ancient samples from Eurasia studied in previous works (The 1000 Genomes Project Consortium, 2015) (Supplementary File 1). The most important contributions to samples included in our data set were 1) 137 ancient individuals in (Olalde *et al.*, 2018) including 72 individuals from Great Britain, 30 from Czech Republic, 24 from Hungary, 14 from Germany and 13 from Russia, 2) 74 ancient individuals in (Mathieson *et al.*, 2015) (31 same samples with 390k in (Haak *et al.*, 2015)), including 49 individuals from Great Britain, 15 from Turkey, 35 from Finland, 8 from Russia, 3) 57 ancient individuals from (Mathieson *et al.*, 2018), including 18 individuals from Great Britain, 11 from Hungary, 7 from Germany, 6 from Finland, 11 from Russia, 5 from Ukraine, 4) 40 ancient individuals in (Lipson *et al.*, 2017), including 6 individuals from Great Britain, 9 from Hungary, 14 from Finland, 4 from Ukraine. For a full list of individuals studied see Table S2. A larger set of genotypes with 5,081 ancient and present-day individuals from the same repository was also considered in analyses. Following the same filtering and imputation procedures as for the first data set, the resulting data contained 123,763 genotypes for 477 present-day European individuals from the 1k Genomes project and 697 ancient samples from previous studies (Supplementary File 2). The data were imputed from genotypes with 20% missing SNPs.

## Supporting information

Supplementary File 1 (Human data)

Supplementary File 2 (Human data)

## Acknowledgements

We thank the Paris-Saclay Center for Data Science 2.0 (IRS) and LRI for funding BD and SL salaries and supporting FJ’s mobility. We also thank Cyril Furtlehner for discussions. This article was developed in the framework of the Grenoble Alpes Data Institute, supported by the French National Research Agency under the “Investissements d’avenir” program (ANR-15-IDEX-02).

## Supplementary Materials

Supplementary tables and figures for “Inference of Population Genetic Structure from Temporal Samples using Bayesian Factor Analysis” by François et al.

**Table S1.**
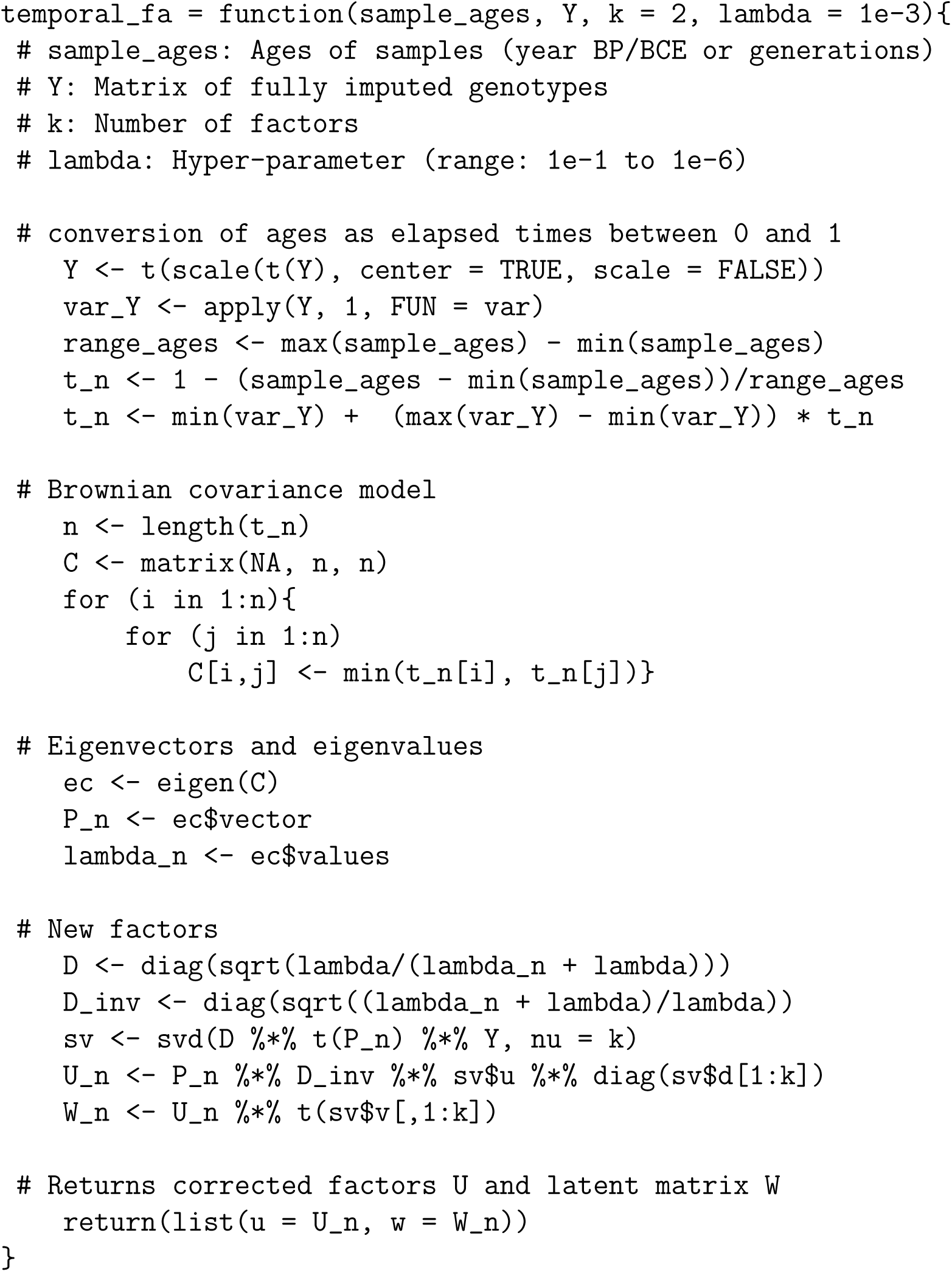
R code for temporal factor analysis

**Figure S1.**
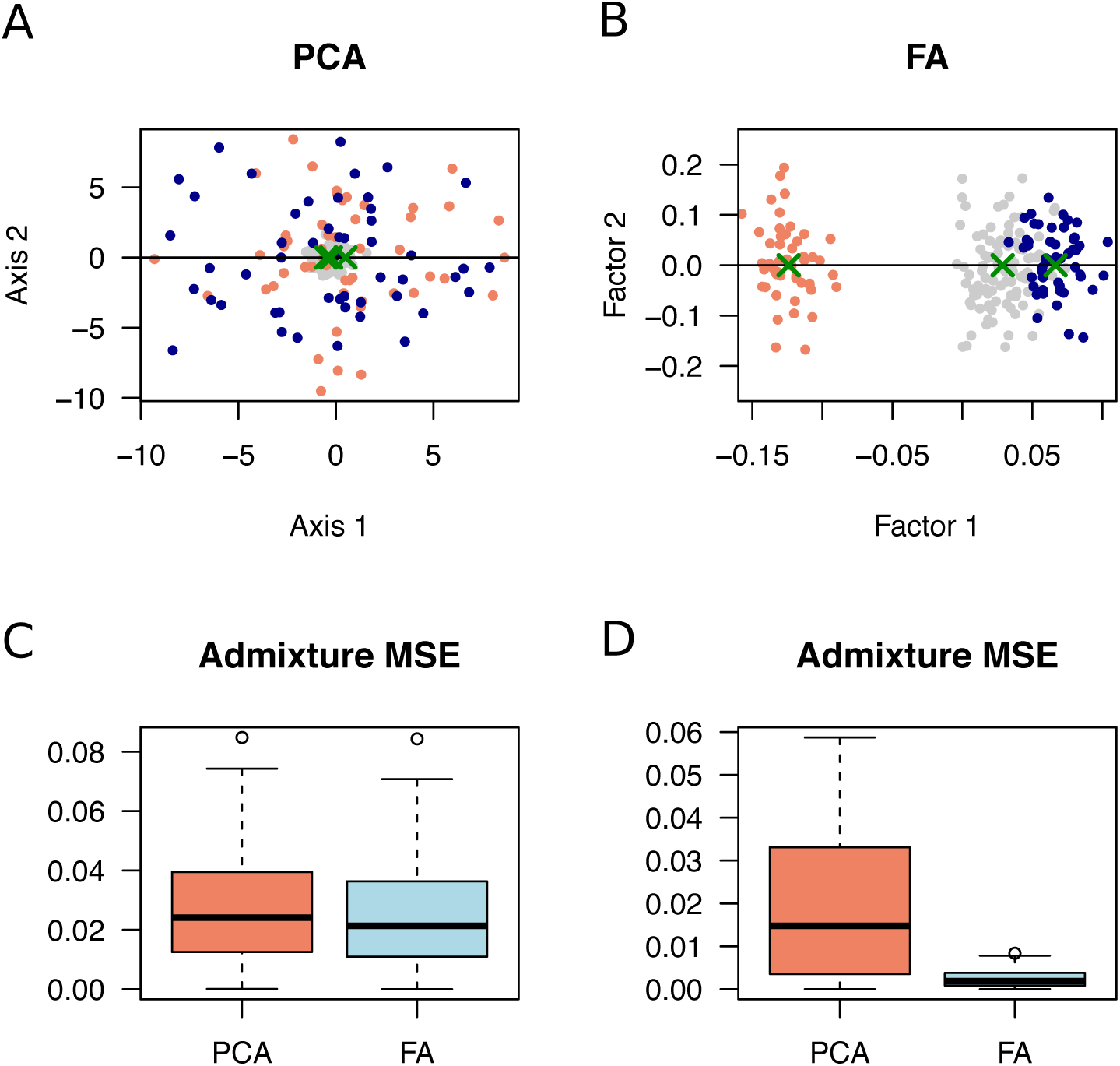
Admixture model simulation. Shrinkage in PC projections. Simulation of two-population admixture models (25-75 % proportions). Two hundred samples with ages equal to 0 (present-day admixed individuals, grey color) and 1,000 generations (ancestors, orange and blue colors) were simulated. A) Plot for PC projection of ancient samples onto the admixed population, with a strong shrinkage effect (coalescent simulation), B) Factor analysis plot showing correction for shrinkage, C) Mean square error for estimates of admixture proportions from PC projections and FA plots (100 generative model simulations), D) Mean square error for estimates of admixture proportions (100 coalescent simulations). Green crosses represent population centers, from which admixture estimates were computed.

**Figure S2.**
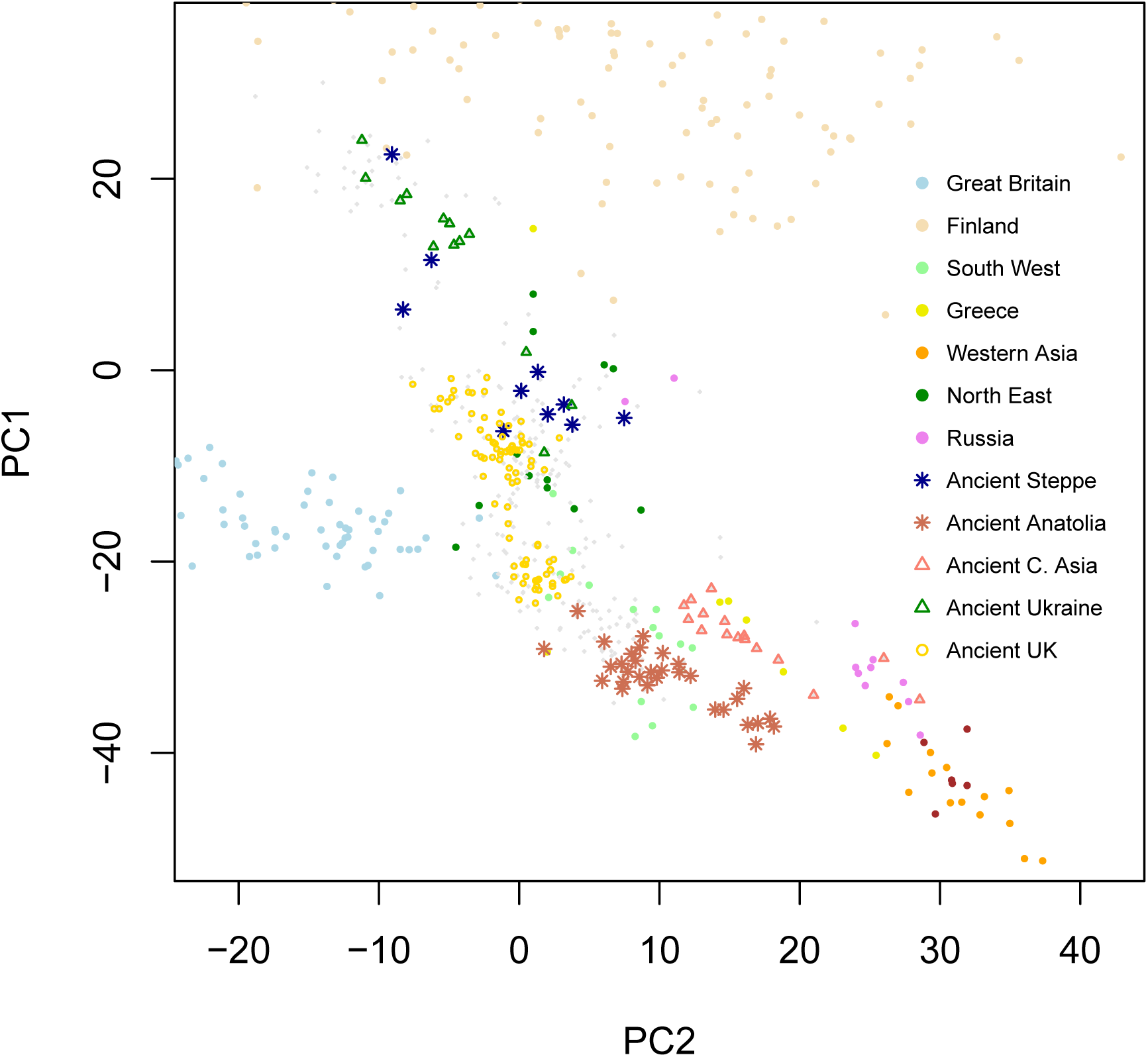
Ancient Humans - PC projections. Projections of 386 ancient Eurasian genomes with age ranging between 400 and 12,000 years BP on principal components of 249 European genomes from the 1,000 Genomes data. Present-day individuals are represented as colored full dots. Smaller light grey dots and other types of dots are ancient genomes.

**Figure S3.**
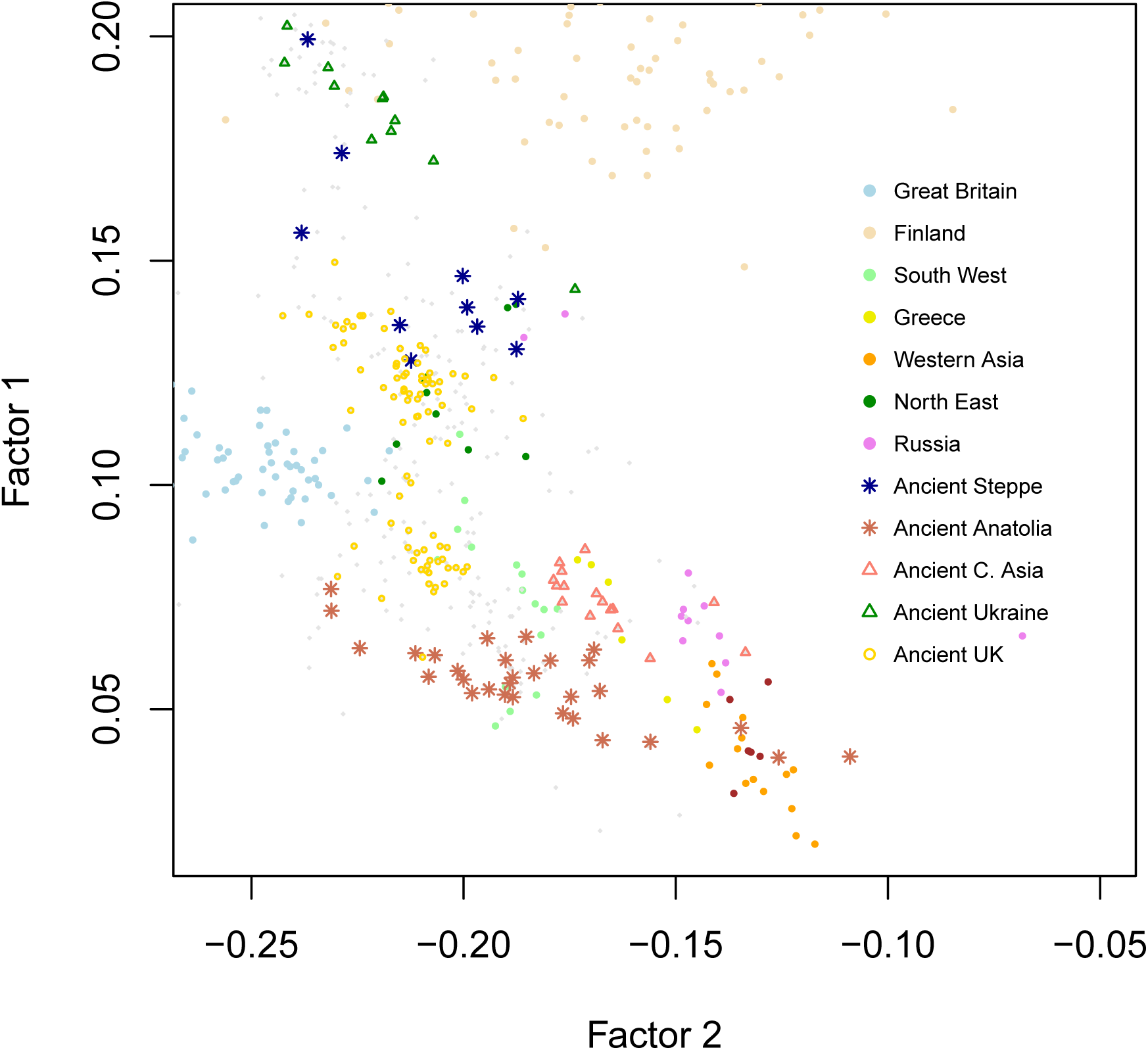
Ancient Humans - Supervised factor analysis. Factor analysis of 386 ancient Eurasian genomes with age ranging between 400 and 12,000 years BP and 249 European genomes from the 1,000 Genomes data. Present-day individuals are represented as colored dots. Smaller light grey dots and other types of dots are ancient genomes.

**Figure S4.**
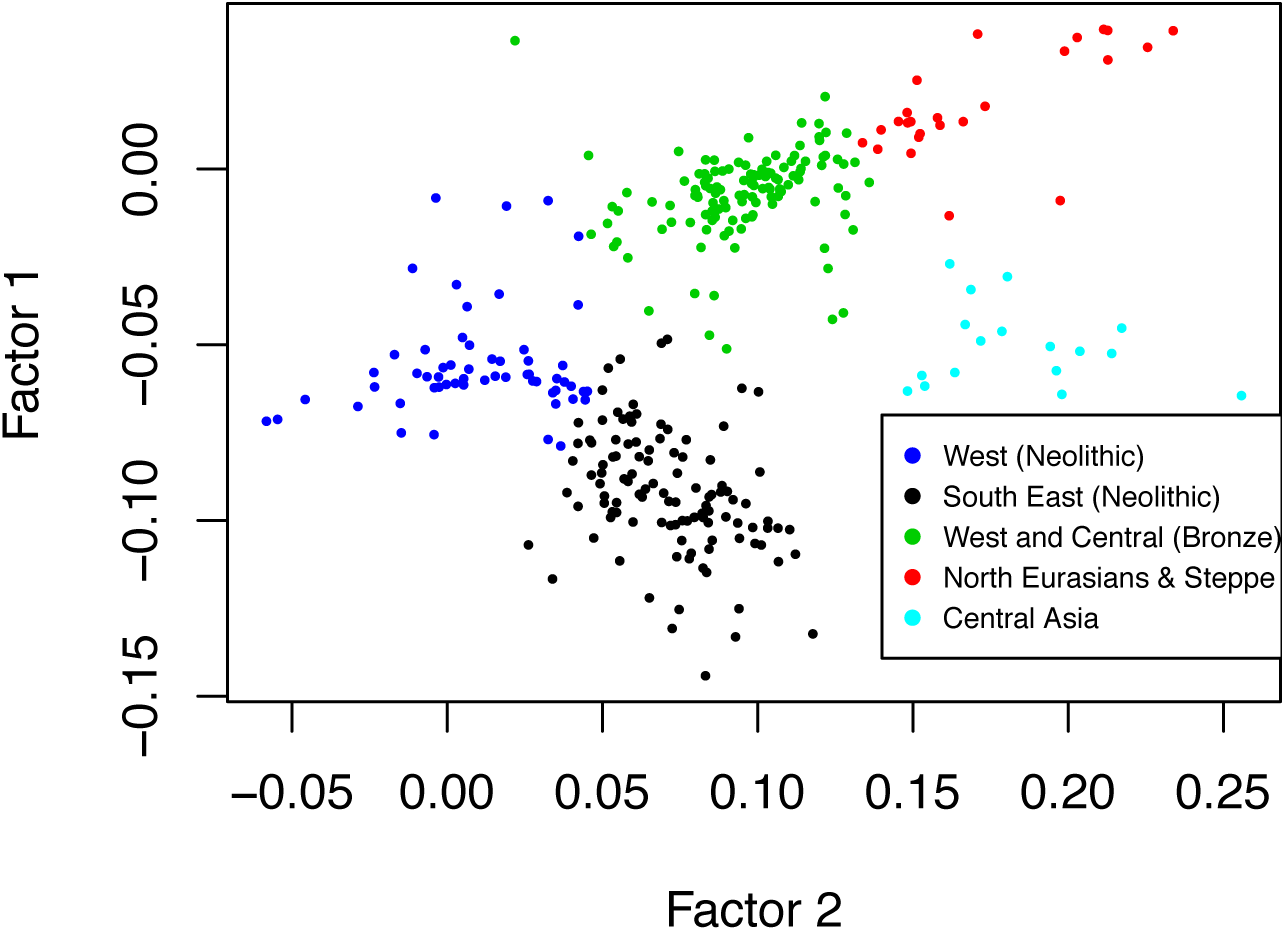
Ancient Europeans - Clusters in factor analysis. Definition of clusters for computing estimates of average ages per factor region. Cluster 1 consists of individuals from Ukraine and Scandinavia, not represented in the plot. Cluster 2 is formed of North Eurasian individuals (Russia, Samara, red color) and central and western Europeans (green color). Cluster 3 is formed of Near Eastern individuals, southern and western Europeans (Neolithic, black and blue colors). Cluster 4 is formed of central Asians (Neolithic, light-blue color). Clustering was performed with a *k*-means algorithm.

**Figure S5.**
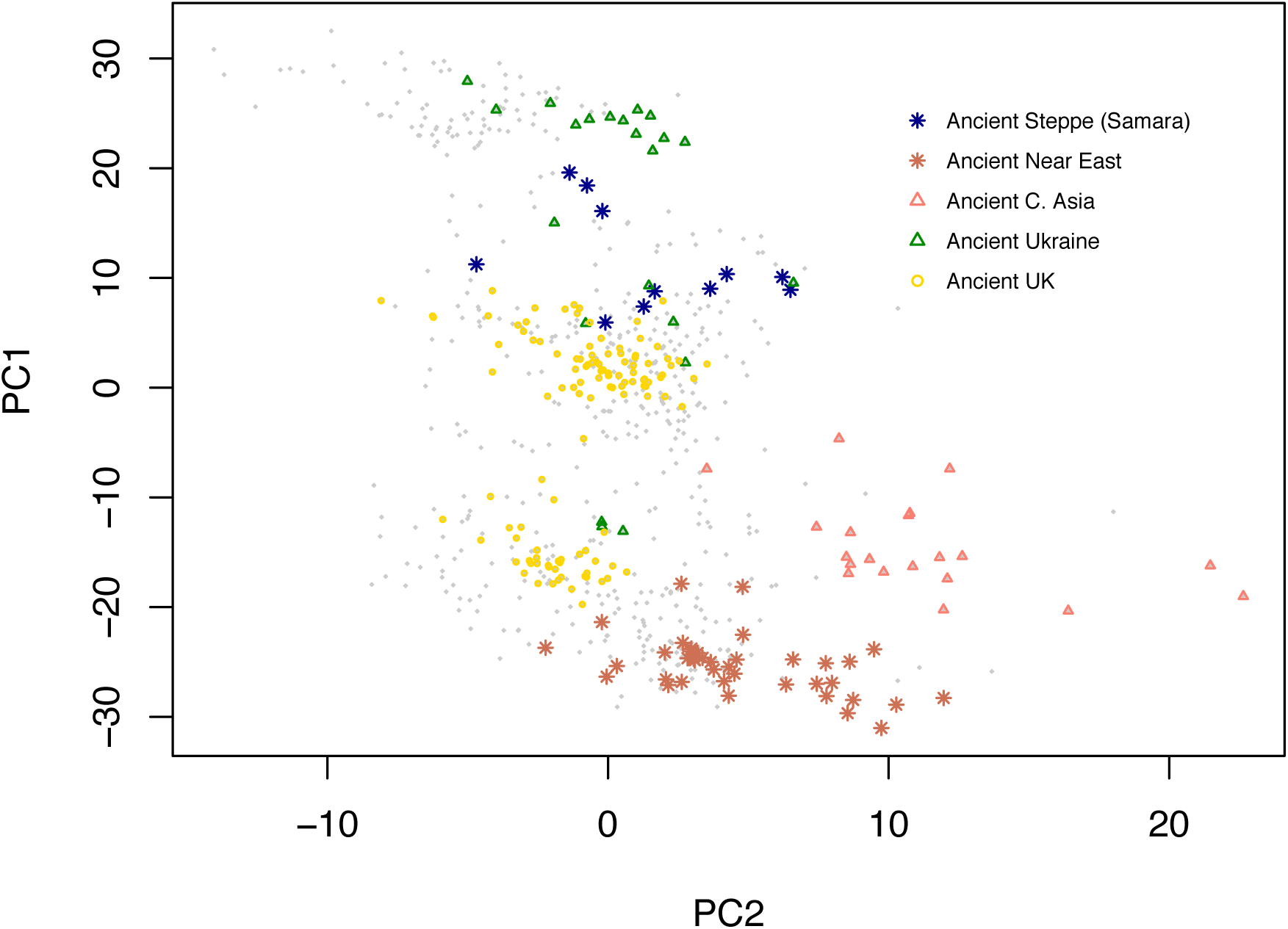
Extended ancient genome data set - PC projections on 1k Genomes data. Projections of 697 ancient genomes on the principal components of 477 genomes from the 1k Genomes data. Only ancient individuals are displayed with some populations emphasized (dates more recent than 12 ky cal BP).

**Figure S6.**
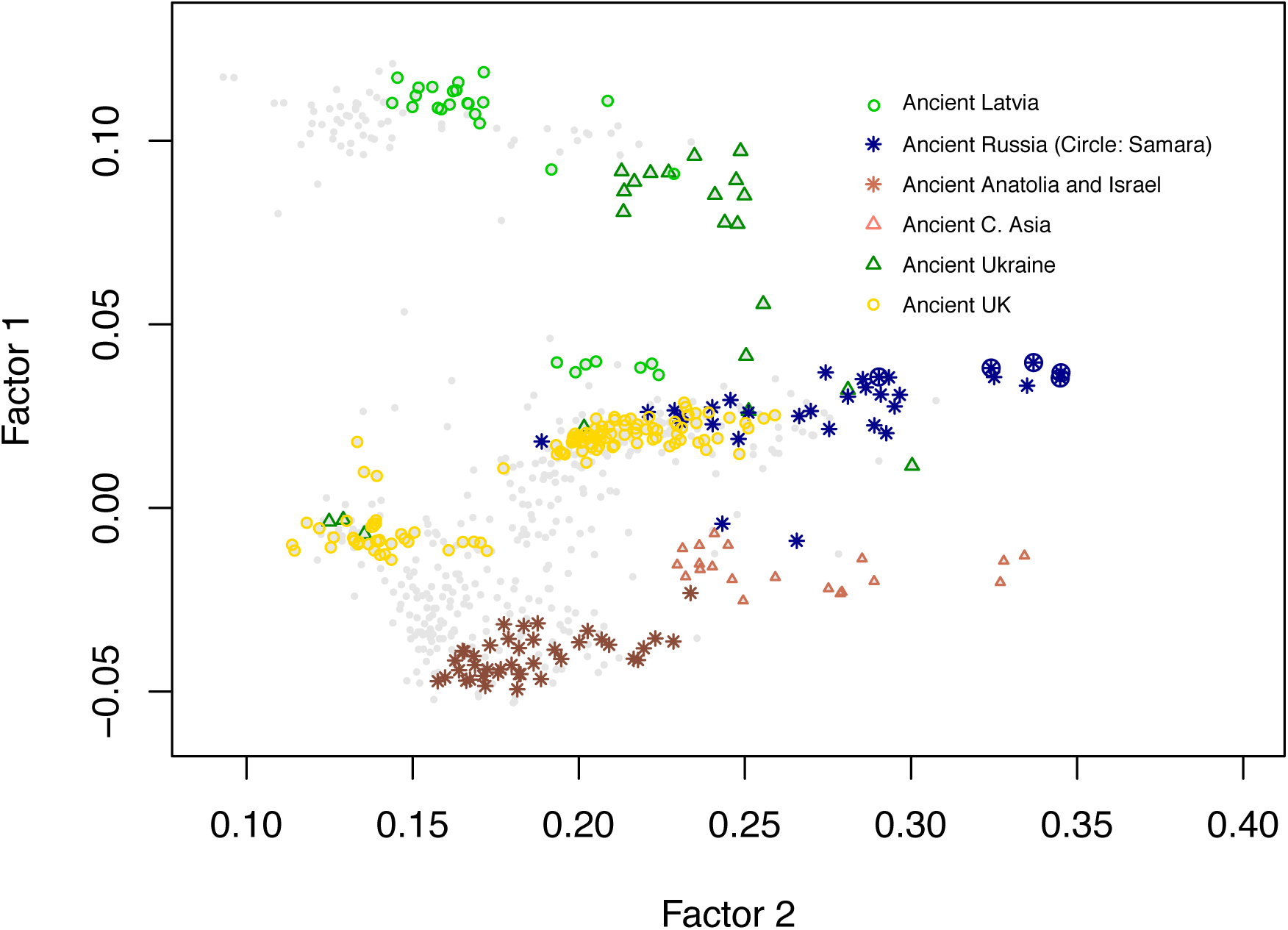
Extended ancient genome data set - Factor Analysis. Factor analysis of 697 ancient genomes with some populations emphasized (dates more recent than 12 ky cal BP). The observed pattern similar to PC projections, but more consistent with geography.

